# DeepSpaceDB: a spatial transcriptomics atlas for interactive in-depth analysis of tissues and tissue microenvironments

**DOI:** 10.1101/2025.01.05.631419

**Authors:** Vladyslav Honcharuk, Afeefa Zainab, Yoshiya Horimoto, Keiko Takemoto, Diego Diez, Shinpei Kawaoka, Alexis Vandenbon

## Abstract

Spatial transcriptomics provides a revolutionary approach to mapping gene expression within tissues, offering critical insights into the spatial organization of cellular and molecular processes. However, generating new spatial transcriptomics data is expensive and technically demanding, and analyzing such data requires advanced bioinformatics expertise. While publicly available datasets are growing rapidly, existing databases offer limited tools for interactive exploration and cross-sample comparisons.

Here, we introduce DeepSpaceDB, a next-generation spatial transcriptomics database designed to address these issues. DeepSpaceDB focuses on interactivity and advanced analytical functionality, enabling users to explore spatial transcriptomics data with unprecedented flexibility. DeepSpaceDB allows for interactive selection and comparison of gene expression across regions within a single tissue slice or between slices, such as comparing hippocampal regions of an Alzheimer’s model mouse and a control. It also includes quality indicators, database-wide trends, and advanced visualizations that provide real-time interactivity, such as zoomable plots and hover-based information display. Moreover, these functions are not restricted to the samples collected in our database but can also be applied to samples uploaded by users.

The current version of DeepSpaceDB focuses explicitly on samples of the 10X Genomics Visium platform, ensuring higher-quality analyses and enhanced exploration tools, including comparison between interactively selected regions of tissue sections. This tradeoff enables unique features like similarity-based sample embeddings and database-wide comparisons, setting it apart from other databases prioritizing broad platform coverage over functionality.

With its combination of advanced tools and interactive capabilities, DeepSpaceDB represents a transformative resource for spatial transcriptomics research, paving the way for deeper insights into tissue organization and disease biology.

**Availability:** DeepSpaceDB is available at www.deepspacedb.com.

## Introduction

Spatial transcriptomics is a revolutionary technology that enables the spatial mapping of gene expression within tissues ^1–3^. The retained positional information of cells is critical for understanding the complex organization of tissues and how cellular behaviors are influenced by their microenvironment. Thus, spatial transcriptomics provides unprecedented insights into the cellular and molecular architecture of biological systems, advancing our understanding of health, disease, and development.

However, generating new samples is financially costly and technically challenging. In addition, analyzing spatial transcriptomics data requires a high level of expertise in bioinformatics. At the same time, publicly available spatial transcriptomics samples are accumulating (see Fig. S1A). There is therefore a demand for databases that allow users to easily and interactively explore existing spatial transcriptomics data. A number of spatial transcriptomics databases have been developed, including SpatialDB, SODB, STOmicsDB and SOAR ^4–7^. Although SpatialDB contains only a limited number of samples and appears to be no longer updated, SODB, SOAR and STOmicsDB contain large numbers of samples covering many different platforms and therefore provide a great service as comprehensive data repositories. However, their use for exploring spatial transcriptomics data is limited because the tools they include are basic and lack interactivity (see Suppl. Table S1). Whereas they allow users to plot the expression of a gene of interest or inspect clustering results, they lack – for example – measures of quality, the ability to freely compare gene expression between arbitrarily selected parts of tissue slices, or the ability for users to upload a sample and compare it to samples in the database. Standalone desktop tools such as Loupe Browser by 10X Genomics allow interactive analysis of single samples, including inspection of quality measures, gene expression and clustering results, but lack the ability to browse through different samples, comparisons between manually selected regions, and inspection of pathway activities.

To address the above weaknesses, we present DeepSpaceDB, a spatial transcriptomics database that allows interactive and smooth exploration of spatial transcriptomics data. The main points of difference between DeepSpaceDB and existing databases are as follows. Compared to existing databases, DeepSpaceDB offers vastly expanded analysis functions with higher interactivity. Of course, DeepSpaceDB includes basic functions such as searching for a specific tissue and condition of interest and visualizing a gene’s expression pattern in a tissue slice. In addition, DeepSpaceDB allows users to interactively select multiple parts within a tissue slice using their mouse cursor and compare gene expression between them. Comparisons can also be made between parts of different tissue slices, such as – for example – between the hippocampal parts of the brain of an Alzheimer’s disease model mouse and that of a healthy control mouse. All plots are interactive and show not merely a static figure but allow users to zoom in and out and show additional information (e.g., gene expression or pathway activities) when hovering above spots with the cursor. In addition, DeepSpaceDB offers database-wide views and the ability to compare samples. This is in contrast with existing databases which treat each sample essentially as an independent and isolated case. For example, quality indicators of samples can easily be compared to the database-wide trends, making it easier for users to judge whether a sample has sufficiently high quality. Furthermore, samples have been processed into 2D embeddings that facilitate the exploration of similar (i.e., nearby samples in the embedding) samples. The above advantages are possible only because we made the tradeoff to limit DeepSpaceDB to samples of only a limited number of transcriptomics platforms; the current version 1.0 includes only samples of the 10X Genomics Visium platform (currently the most widely used platform). In the future, samples of other popular platforms will be added. Note that this approach is the opposite of other spatial transcriptomics databases, which put more emphasis on wide coverage but with limited analysis and exploration functions. Furthermore, DeepSpaceDB allows users to upload their own samples and apply many of the functions mentioned above on their own data, without the need for programming or data analysis experience.

With its unique combination of advanced functionality and user interactivity, DeepSpaceDB sets a new standard for spatial transcriptomics databases. As we expand to include more platforms, DeepSpaceDB is poised to become the go-to resource for the spatial biology community.

## Results

### An overview of collected samples

We collected Visium samples from NCBI GEO, the 10X Genomics website, and other data repositories (see Suppl. Table S2), and processed them through a pipeline that included initial processing, quality checks, normalization, and downstream analyses (Figure 1A, see Methods for details). Initial exploratory analysis of the collected data included a quality assessment. We observed a relationship between UMI counts and detected genes, similar to the tendency that is often observed for single-cell data (Fig. S1B,C). However, it has been pointed out that these quality indicators at least partly reflect biological properties of the tissue of origin ^8^. Here, too, we observed clear differences in the number of detected genes between different tissues (Fig. S1D,E). In addition, we observed a strong association between these quality measures of each spot and its number of neighboring spots (Fig. S1F-I). We labeled 38,484 spots (out of 3.9 million spots; 0.97%) with low numbers of detected genes and isolated spots (i.e. spots lacking direct neighboring spots) as having low quality. After filtering out samples with >25% low quality spots, a total of 1,674 samples (1,011 human and 663 mouse) Visium samples remained. Frequent tissues of origin included lung, brain, skin, and breast for human, and brain and liver for mouse (Fig. 1B,C). Many human samples were related to diseases, especially cancer (Fig. 1B), and healthy human samples were rare. However, human “healthy” samples are likely to also include samples that simply lack a pathological annotation. Mouse samples contained more healthy control samples (Fig. 1C). In a second step of exploratory analysis, we processed each sample into a pseudo-bulk sample by averaging gene expression across all spatial locations, and processed these into a 2D embedding (human: Fig. 1D, mouse: Suppl. Fig. S2A). We observed that, in general, samples from the same or related tissues tend to group together (i.e., have relatively similar gene expression patterns). However, especially for human samples there is also a prominent group of samples originating from cancer patients (Fig. 1D, samples marked by dotted line; Suppl. Fig. S2B), where the grouping per tissue of origin is obfuscated.

**Figure 1:**
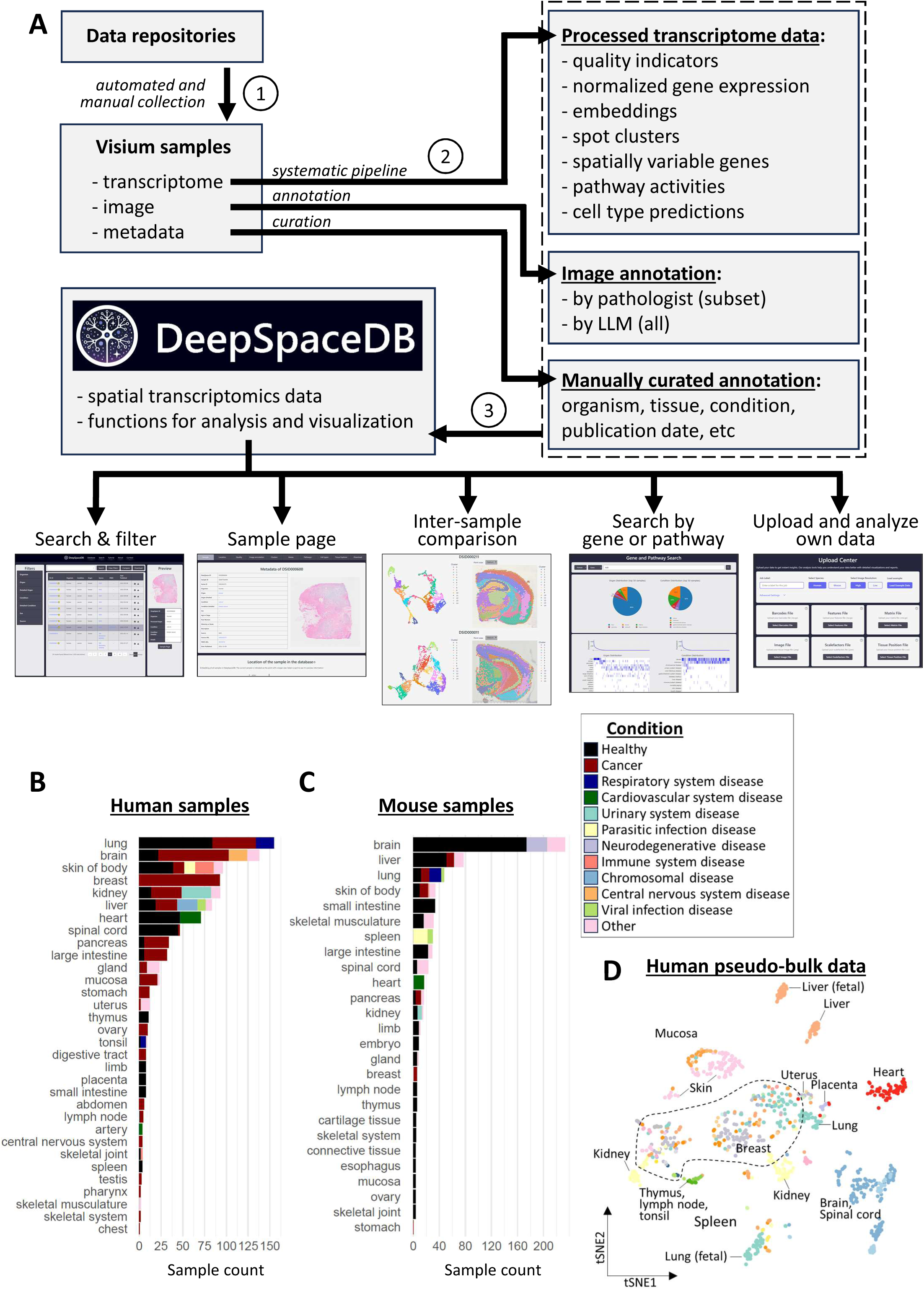
Spatial transcriptomics data collection of DeepSpaceDB **(A)** Schematic overview of the data collection and processing workflow. First, transcriptomics, image and annotation data were collected from various sources. Next, each data type was processed and analyzed. Finally, resulting data was included in DeepSpaceDB along with functions for analysis and visualization. **(B)** The number of human samples per tissue of origin. Colors indicate the condition. **(C)** Same as (B) for mouse samples. **(D)** Embedding (tSNE plot) of human samples after processing to pseudobulk data. Colors indicate the tissue of origin. Prominent tissues are indicated. The dotted area indicates a large group of cancer-related samples (see also Suppl. Fig. S2B).

We similarly explored the data on the level of individual spots. After clustering all human and mouse spots by similarity of gene expression into 50 clusters (see Methods; Suppl. Fig. S2C,D), we attempted to assign a rough annotation to each cluster of spots based on their association with sample tissue and condition annotations, and the gene expression patterns within each cluster (Suppl. Fig. S3 for human and Suppl. Fig. S4 for mouse). Note that this annotation process is complicated by 1) the fact that spots reflect the average gene expression of multiple cells resulting in few clearly distinct clusters, and 2) the lack of spot-level annotations (e.g., not all spots in a cancer-related sample are necessarily cancer tissue). Nevertheless, we could assign an annotation to most of the spot clusters, and inspect the distribution of clusters with common and different annotation within the 2D embedding. For example, for human, clusters associated with tumor tissue, neurons, tissue enriched with immune cells and liver fill up distinct subspaces in the 2D embedding (Suppl. Fig. S5). Similar trends are seen for clusters associated with brain tissue, neurons (brain and spinal cord), liver, and immune-enriched tissue in mouse (Suppl. Fig. S6).

Below follows a tour of the DeepSpaceDB database, including the exploration of a sample of interest, comparing between pairs of samples, and searching the database with a query gene or pathway of interest.

### Exploring a sample

Under the Database tab, users can search the collection of samples for their species, tissue, condition and other characteristics of interest. Selecting a sample shows a preview and takes the user to a sample-oriented page. Below, we will walk through this page and introduce its features. For this we will use a human breast cancer sample (DeepSpaceDB internal ID DSID000600), explore its features and use it to compare between tumor and non-tumor tissue in this sample ^9^.

#### Annotation data

This shows background of the sample, including the data source (such as NCBI GEO, the 10X website, etc), the tissue and condition (based on UBERON and DO or similar ontologies), and where available the sex, age, strain or ethnicity of the patient or animal, and links to supporting scientific literature. A short description of the sample is also provided.

#### Location of the sample in the database

One of the weak points of existing spatial omics databases is that samples are essentially treated and presented in isolation of each other, without integrating them into a database that allows inter-sample comparison. In DeepSpaceDB, however, all collected data has been integrated, allowing easier comparison between them. In this section, a 2D embedding is shown of all samples in the database, indicating the currently selected sample. This allows the user to quickly find similar samples in the database. Here, for the currently selected breast cancer sample, we can see that similar samples include breast cancer samples, but also clear cell renal carcinoma, and ovarian and lung cancer samples (Fig. 2A).

**Figure 2:**
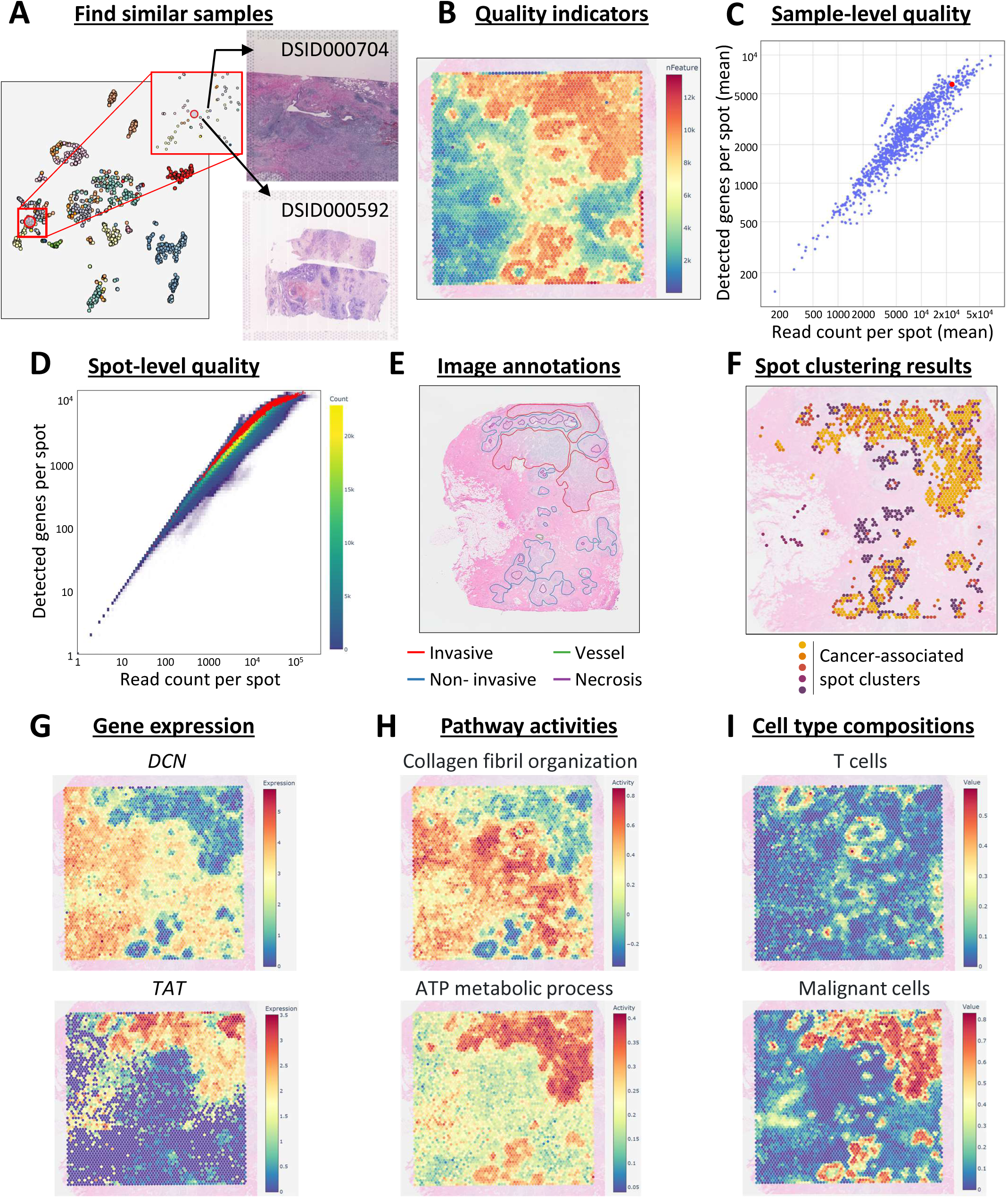
Inspection of a human breast cancer sample in DeepSpaceDB **(A)** Using the 2D embedding (tSNE plot) to inspect samples that are similar to the currently selected sample. The images of two similar samples are shown as example. **(B)** Plot of the number of detected genes at each location inside the tissue slice. **(C)** The average number of reads (X axis) and the average number of detected genes (Y axis) for all human samples in DeepSpaceDB. The currently selected sample is indicated in red. It has a relatively high number of reads and detected genes compared to other samples. **(D)** The number of reads (X axis) and detected genes (Y axis) per spot in the database. Spots of the currently selected sample are indicated in red. **(E)** The H&E image of the current sample with annotations by a human expert. **(F)** Spots in the current sample which were assigned to spot clusters associated with tumor tissue. Spots assigned to other clusters are not shown. **(G)** Examples of high-scoring SVGs: *DCN* and *TAT*. **(H)** Examples of high-scoring spatially variable biological processes. **(I)** Prediction of the distribution of T cells (top) and malignant cells (bottom) in the tissue slice.

#### Quality measures

It is often critical to confirm if a sample is of sufficiently high quality compared to other samples before using it to generate hypotheses. DeepSpaceDB offers various ways to check the quality of a sample and the spots it contains. The measures include the number of detected genes, the number of UMIs, the percentage of reads originating from mitochondrial genes, and the number of neighboring spots of each spot. These quality measures can be visualized within the tissue slice (Fig. 2B). In addition, the quality of a sample can easily be compared with all other samples in the database, a feature which is critically missing from exiting databases. For example, we can see that the currently selected sample has a relatively high average number of reads per spot and average number of detected genes per spot as compared to other samples in the database (Fig. 2C). The same comparison can be made on the level of individual spots (Fig. 2D).

#### Image annotations

Image annotations can assist the interpretation of spatial patterns inside tissues. In DeepSpaceDB (version 1.0), 69 human breast cancers have been manually annotated by a pathologist specializing in breast pathology. In the current sample, several invasive and non-invasive tumors were annotated, in addition to other features such as normal ducts and vessels (Fig. 2E). However, the manual annotation by a pathologist takes time and effort, and each pathologist tends to be specialized in one specific tissue. To collect manual annotations for all images in DeepSpaceDB is therefore a daunting task. Moreover, the Visium images are typically of lower resolution than typical images used for diagnosis. As an alternative, we therefore used an LLM to describe features inside each image, using a 5-by-5 grid. Although the promise of AI-assisted annotation of histopathology images has been demonstrated ^10^, and these predictions can help interpreting the spatial transcriptomics patterns, they need to be treated with care.

#### Spot clustering

Another way to assist the interpretation of spatial patterns of gene expression is to cluster spots by the similarity of gene expression patterns. In DeepSpaceDB, spots of each sample were clustered in 2 ways: 1) within-sample clustering, and 2) global clustering across the entire database (see Methods, and Suppl. Fig. S3,4). Both clustering results can easily be visualized. We assigned annotations to the global clusters, which can assist in the exploration of structures and features within the tissue section. For example, for the selected breast cancer sample, we can see that spots assigned to tumor-associated clusters are accumulated at specific locations within the section (Fig. 2F).

#### Spatially variable genes and biological processes

One advantage of spatial transcriptomics data is that it enables the visualization of gene expression within a tissue. In many cases, a subset of genes has a clearly distinct spatial expression pattern, reflecting substructures within the tissue. In DeepSpaceDB, SVGs in each sample have been precalculated by the singleCellHaystack method ^11,12^. Users can search and select genes of interest, and instantly and interactively plot their expression patterns within the tissue slices along with the tissue image. In addition to individual genes, in DeepSpaceDB, the activity of sets of genes involved in a common biological process can be easily visualized (see Methods). We used singleCellHaystack to predict the significance of the spatial pattern of activity of the biological processes in a similar way as we did for individual genes. As an example, in the selected breast cancer sample, top scoring SVGs follow distinct patterns, exemplified by the genes decorin (*DCN*) and tyrosine aminotransferase *(TAT*) (Fig. 2G). Two of the high-scoring biological pathways in this sample are collagen fibril organization and ATP metabolic process, which have high activity in the non-cancerous and invasive tumor spots, respectively (Fig. 2H).

#### Cell type composition

For each sample in DeepSpaceDB, the cell type compositions at each spatial location were predicted using the method RCTD (see Methods) ^13^. In brief, RCTD uses a reference scRNA-seq dataset to deconvolve the gene expression pattern of each Visium spot. Fig. 2I shows how RCTD predicted the presence of T cells and malignant cells at specific locations within the tissue slice, fitting with the observations described above.

### Interactive exploration of gene expression patterns

The above exploration suggests that the currently selected sample contains invasive tumor tissue at the top of the slice, and a relatively smaller amount of non-invasive tumor tissue at the bottom. We were interested in obtaining hints as to the difference between those two regions, and the region located between them. DeepSpaceDB allows users to freely and interactively (using the mouse cursor) select multiple parts of a tissue slice and compare gene expression patterns between the selected parts. As an example, here we selected three parts within the tissue: the region at the bottom covering the smaller tumor tissue (“set 1”, 418 spots), the region at the top covering the larger tumor tissue (“set 2”, 1,058 spots), and the region between the tumor tissue (“set 3”, 461 spots) (Fig. 3A). DeepSpaceDB calculates the average gene expression in each of the selected parts and returns the results in the form of a table and scatter plots (Fig. 3B). Here, for example, we can see that the genes *CDH2* and *CEACAM6* have relatively high expression in the larger tumor tissue at the top and the smaller tumor tissue at the bottom of the slice, respectively (Fig. 3C). Similarly, we can easily find and inspect genes with large differences in expression between sets 1-3 and between sets 2-3 (Suppl. Fig. S7). In addition to manually selected sets of spots, comparisons between spot clusters are also possible.

**Figure 3:**
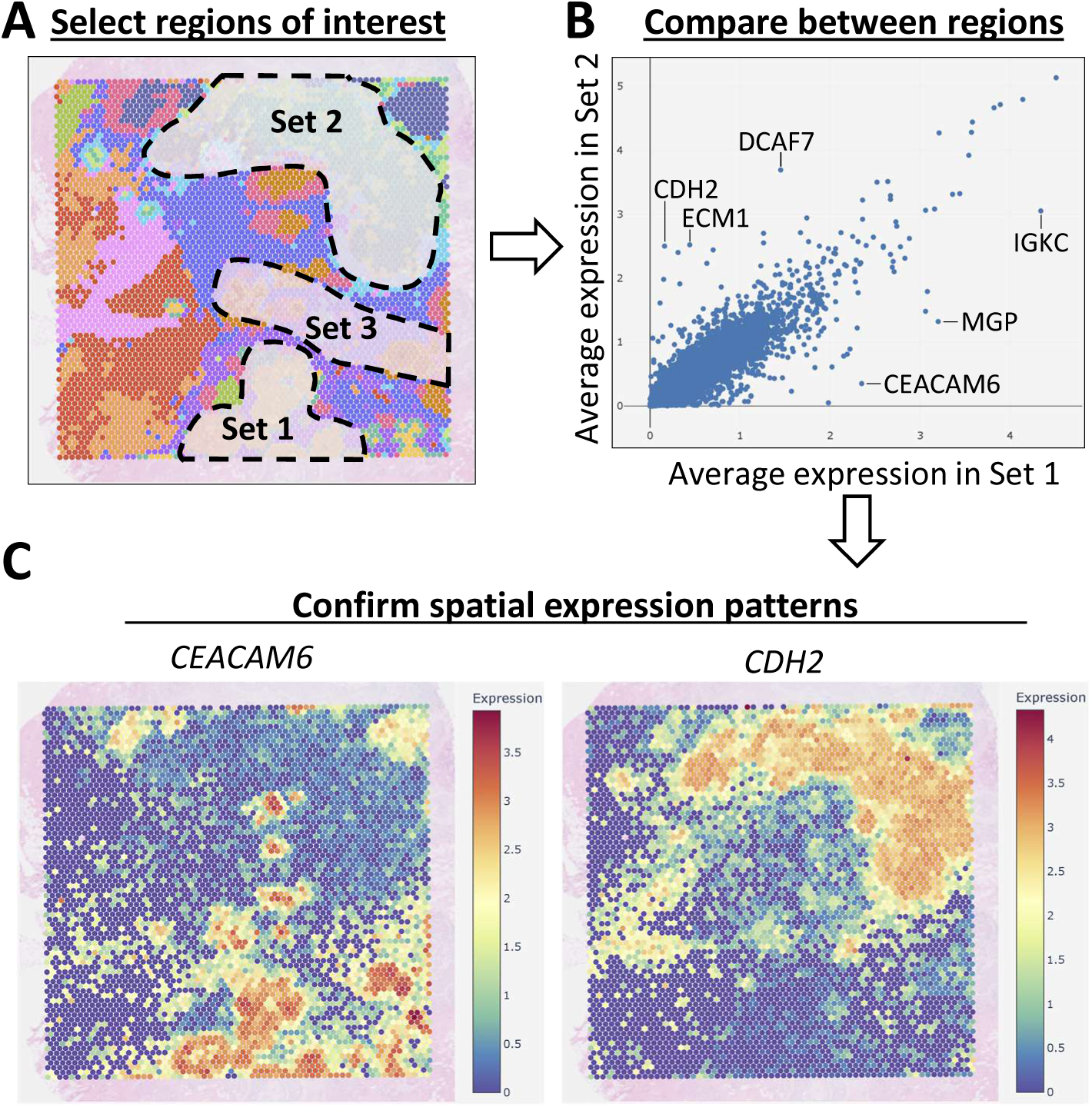
Comparing gene expression between different regions within a sample **(A)** The three selected regions are indicated. Selections are made using the mouse cursor. **(B)** Scatterplot of the average gene expression in set 1 (X axis) and set 2 (Y axis). A number of genes with large differences is indicated. Similar comparisons between sets 1 and 3, and between sets 2 and 3 are shown in Suppl. Fig. S7. **(C)** The spatial expression patterns of two selected genes are shown. *CEACAM6* has a higher expression in set 1 than in set 2. *CDH2* has a higher expression in set 2 than in set 1.

### Download data

At the bottom of the sample page, DeepSpaceDB allows users to download various data, including the image and transcriptomics data of the sample. We also provide data files of pre-calculated features, such as the predicted spatially variable genes and biological processes predicted by singleCellHaystack, and Seurat objects. Functions for downloading data of multiple samples are also available on the “Database” tab.

### Comparison between different tissue slices

In addition to comparing gene expression between different parts of a single tissue, DeepSpaceDB also allows in-depth comparisons between two different tissues. Here we describe a comparison between the hippocampal part of an Alzheimer’s disease mouse model brain (sample ID DSID000211) and a healthy control brain (sample ID DSID000011) ^14,15^. Figure 4A summarizes the interactive selection using the cursor of the hippocampus of both samples. The result of the comparison is a table with the average expression of all genes in both selections and a scatterplot summarizing the comparison (Fig. 4B). Inspection of the expression of genes *Bc1*, *Prnp*, and *Atp6v0c* confirms their higher expression in the hippocampus (and other regions) of the Alzheimer’s disease model brain (Fig. 4C).

**Figure 4:**
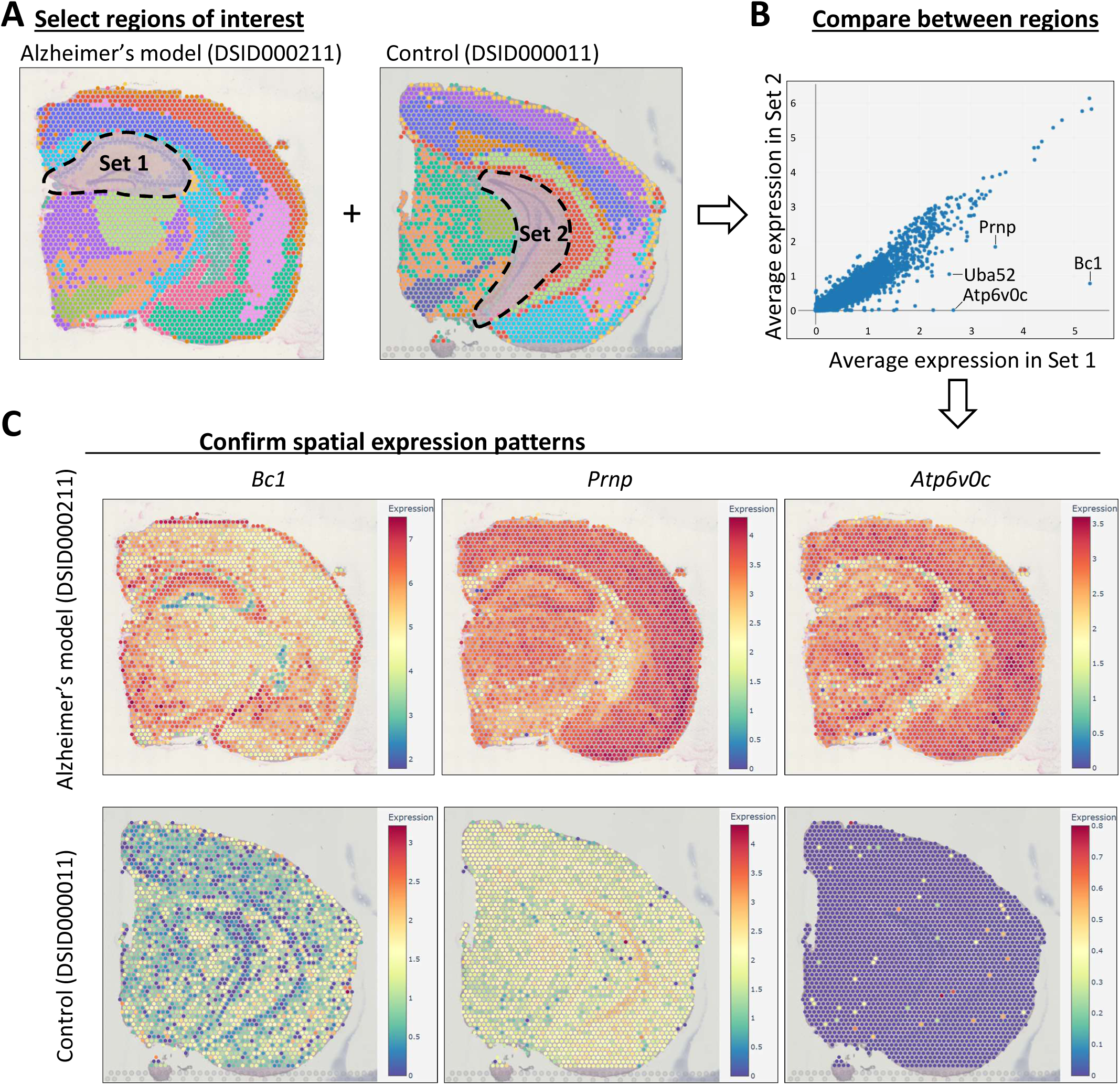
Comparing gene expression between different samples **(A)** The selected regions in an Alzheimer’s disease brain model (left) and a control brain (right) is shown. **(B)** Scatterplot of the average gene expression in the sets of selected spots of the two samples. A number of genes with high expression in the Alzheimer’s disease model sample is indicated. **(C)** The spatial expression patterns of three selected genes are shown in the Alzheimer’s disease model samples (top) and in the control sample (bottom).

### Searching DeepSpaceDB using a query gene or biological process

The above examples illustrate how to browse DeepSpaceDB starting from a tissue or condition of interest. An alternative is to search DeepSpaceDB using a query gene of interest, similar to the way biologists might search for a gene in literature or in a genome browser. As a result, DeepSpaceDB returns all samples in the database, ordered by the degree to which the query gene is a SVG as judged by the singleCellHaystack method. This allows users to find samples in which their gene of interest has varied expression within the tissue. Moreover, the ordering of samples and the tissue (or condition) they originated from can give valuable insights about the function of the query gene. In addition to searching using query genes, users can also use biological pathways as queries. Several examples are shown in Fig. 5 (left: summary of the sample ordering, right: a high-scoring sample). For the mouse *Ttr* gene, top-ranking samples are dominated by brain samples (Fig. 5A, left), such as sample DSID000639, where *Ttr* has highly concentrated expression in two locations within the brain (Fig. 5A, right). For the human *IL7R* gene, there is no clear tendency for a tissue of origin (Fig. 5B, left). One of the top-scoring samples is a tonsil sample, in which *IL7R* shows a complex expression pattern across the tissue slice (Fig. 5B, right). For the xenobiotic metabolic process, high-scoring samples tend to include lung and liver, but one of the top samples is a kidney sample where this pathway is more active in the renal cortex than in the renal medulla (Fig. 5C). For the phagocytosis recognition biological process, there is no strong tendency among tissues of origin (not shown), but cancer-related samples tend to be among the high-scoring samples (Fig. 5D, left), possibly reflecting the activation of phagocytosis-related genes in these samples. One example is a colorectal cancer sample (Fig. 5D, right).

**Figure 5:**
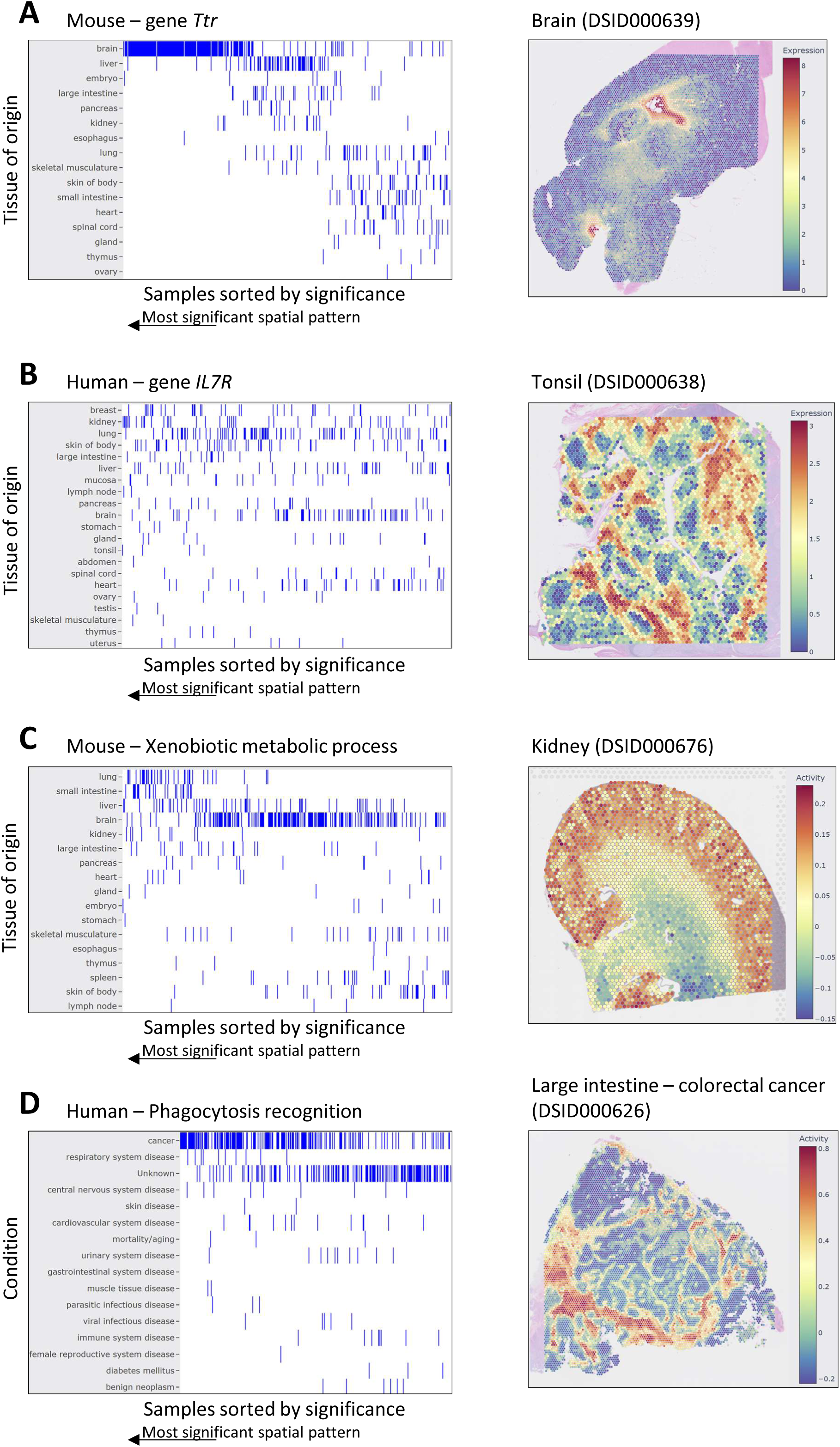
Searching DeepSpaceDB with a query gene or pathway For each query a ranking of all samples sorted by significance of the spatial pattern of the query gene/pathway is shown on the left. Significant spatial patterns here refers to highly non-uniform spatial patterns. Tissues of origin or conditions are indicated. On the right, one of the top 3 high-scoring samples (i.e., samples with strong spatial patterns for the gene or pathway in question) is shown with the expression/activity of the query gene/pathway. **(A)** Searching the mouse samples using query gene *Ttr*. High-scoring samples are strongly enriched in brain samples. **(B)** Searching the human samples using query gene *IL7R*. **(C)** Searching the mouse samples using query Xenobiotic metabolic pathway. High-scoring samples include lung, small intestine, and liver. **(D)** Searching the mouse samples using query pathway phagocytosis recognition. High-scoring samples are biased towards cancer-related samples.

### Analysis of uploaded samples

Users can upload Visium samples to DeepSpaceDB (see Supplementary Fig. S8 for a summary of the sample upload page). After uploading the necessary input files, samples are processed in the same way as the public data included in our database, including normalization, calculation of quality indicators, dimensionality reduction, clustering, and prediction of SVGs. Moreover, users can compare the uploaded samples to samples in DeepSpaceDB. Uploaded data will be stored on the DeepSpaceDB server for a limited time after which it is automatically removed. Each uploaded sample can be accessed through a unique URL which can be shared with collaborators, but data uploaded by users is not incorporated into the DeepSpaceDB data and is not made public to other users.

## Discussion

We present DeepSpaceDB, an integrated database for the interactive exploration and analysis of spatial transcriptomics data. Whereas existing databases focus on the collection of samples of many different platforms, DeepSpaceDB focuses on the implementation of tools that allow the user to explore spatial data more deeply, interactively and intuitively. To make this possible, we restricted the current version of DeepSpaceDB to data of the Visium platform, which has both a genome-wide coverage and many available samples. However, the field of spatial transcriptomics is still in rapid development, and it is reasonable to expect new platforms will be introduced in the next few years. We will monitor further developments and expand the database to cover future popular platforms, such as Xenium, Stereo-seq, or other platforms still under development. Undoubtedly, novel strategies will be needed to process the data of these higher-resolution platforms into formats that ensure a smooth experience for users.

DeepSpaceDB allows users to easily inspect the quality of samples (e.g., number of detected genes, read counts, etc) and compare it with other samples, offers integrated views of the location of each sample w.r.t. other samples, enabling users to easily find similar (or different) tissue slices. DeepSpaceDB also provides several levels of sub-tissue annotation: 1) spots have been grouped into database-wide clusters to which we have assigned annotations, 2) an LLM was used to detect structural and pathological features in the image data of each sample, and 3) high-quality annotations by a histology expert are available for a limited but increasing number of samples. Such annotations are absent in existing databases. Spatially variable genes have been pre-calculated in each sample, allowing for a smooth exploration by the user, without having to wait for each plot to be generated. Whereas the plotting of single genes is possible in existing databases, DeepSpaceDB also contains precalculated estimates of biological process activities in each sample. Here too, the statistical significance of spatial patterns of activity has been precalculated, allowing quick and easy visualization. Given the noisy nature of individual genes in spatial transcriptomics data, these pathway activities offer an additional tool for exploration to the users. Cell type predictions within each sample are based on curated reference single-cell datasets. Users are also able to interactively and freely select parts of tissue slices and compare gene expression patterns between these selected parts of interest. This can provide valuable hints about the difference in expression between – for example – tumors versus their immediate surroundings, between different parts of the brain, or between two different germinal centers inside a lymph node. Moreover, comparisons are also possible between different tissue slices. Users therefore can – for example – manually select the same part of the brain of a healthy control mouse and an Alzheimer’s disease model mouse and compare gene expression between these parts. Moreover, many of the above tools can also be applied on samples uploaded by the users. DeepSpaceDB can also be searched to find tissue sections that have a strong spatial pattern of expression for a given query gene of interest. The above functions allow users to interactively explore spatial transcriptomics data in a way that would be impossible to do in existing databases. Finally, raw and processed data as well as pre-calculated analysis results files are available for download for each sample, facilitating further downstream analysis by users.

## Materials and methods

### Data collection and quality control

Transcriptome, image and meta data was collected from the data sources (see Suppl. Table S2), Each sample was manually assigned a tissue of origin (using the Uberon anatomy ontology) ^16^, condition information (Human Disease Ontology, DO) ^17^, as well as information related to the age or state of development, ethnicity or mouse strain, a link to scientific publications, and publication date, where possible.

Data of each sample was processed using the R Seurat package (version 5.0.1) ^18^. Quality control included calculation of the number of detected genes and the UMI counts. After inspection (see Results), we employed a relatively loose quality filter, marking 38,484 spots (out of 3,947,288 spots, 0.97%) with no direct neighboring spots and spots with less than 100 detected genes as having low quality. Samples with >25% low quality spots were filtered out. After removal of low-quality samples, a total of 1,674 samples remained (1,011 human and 663 mouse).

### Transcriptome data processing

To allow for easier inter-sample and inter-location comparison, we integrated the collected data on two levels: the level of spots and the levels of samples (pseudo-bulk). For the sample level, we calculated the average gene expression for each sample across all its spatial locations, transforming it into a pseudo-bulk sample. We converted these pseudo-bulk samples into a Seurat object. This Seurat object was processed in a similar way as a typical a single-cell dataset, including scaling, selecting of highly variable genes (from the set of genes common to all samples), and dimensionality reduction using PCA and tSNE (using 10 principal components; see Fig. 1D and Fig. S2A). Default parameters were used. We performed the spot-level integration in a similar way, including a batch effect correction step using Harmony (version 1.2.0) using the first 50 principal components ^19^, followed by dimensionality reduction using tSNE using the first 50 dimensions returned by Harmony, resulting in 2D tSNE embeddings of spots (Fig. S2C,D). To further annotate the variety of spots in the collection, we clustered spots into 50 clusters using k-means clustering and gave these clusters an annotation after inspection of the properties of spots they contain (tissue of origin, conditions, gene expression; Suppl. Fig. S3,4). The overlap between clusters of spots and tissue and condition annotations was evaluated using the Jaccard index between each cluster and each annotation using the R package scmisc (version 0.8.4) ^20^. Jaccard index values were converted into Z scores using the means and standard deviations of Jaccard index values of 100 random permutated datasets.

In each Visium sample, spatially variable genes (SVGs) were predicted using the singleCellHaystack (version 1.0.2) method ^11,12^. In brief, singleCellHaystack predicts features (genes) with non-random patterns of activity (gene expression levels) inside an input space. Here, we used the 2D spatial coordinates of the spots of each sample as input space to singleCellHaystack. The grid.points parameter of singleCellHaystack was set to a relatively high value (the number of spots divided by 7) but singleCellHaystack was otherwise used with default parameters.

The activities of biological processes were estimated using “module scores” as previously described ^21^. In brief, we used the R package msigdbr (version 7.5.1) to define sets of genes associated with Gene Ontology (GO) biological processes ^22^. For these gene sets, we calculated module scores using Seurat’s AddModuleScore function in each spot of all Visium samples, using default parameters. We predicted spatially variable pathways using singleCellHaystack, in the same way as we did for predicting SVGs, using the module scores of pathways as input.

For cell type deconvolution we used robust cell type decomposition (RCTD) ^13^. We collected single-cell RNA-seq (scRNA-seq) reference datasets from CELLxGENE such that for each Visium sample we had a corresponding scRNA-seq dataset originating from the same (or a similar) tissue ^23^. We limited reference datasets to data sequenced using 10x Genomics platforms, to limit technical differences between Visium data and reference datasets. Many Visium samples are affected by medical conditions, such as cancer or immune diseases, and cell types and states in the reference datasets are likely to not match perfectly with the cell types and states in the tissue slices. Improving the match between references and spatial transcriptomics datasets is one of our long-term goals. Here, for the current version of DeepSpaceDB, we limited ourselves to adding cancer cells to reference datasets. To do so, we first downloaded the tumor scRNA-seq data from the Tumor Immune Single-cell Hub 2 (TISCH2) database ^24^. We then extracted malignant cells from these tumor tissue datasets and added these cells to the corresponding tissue scRNA-seq reference datasets, and used these augmented datasets as reference for cell type deconvolution of Visium samples that were affected by cancer. Where needed, Ensembl IDs were converted to gene symbols using function mapIds of the R package AnnotationDbi (version 1.64.1) ^25^. Furthermore, cell types for which there were less than 25 cells in a reference dataset were excluded, as RCTD requires at least 25 cells per cell type annotation. RCTD was run using doublet_mode = “full” and using otherwise default parameters.

### Image data processing and annotation

Using NDP.view 2.9.29 (Hamamatsu Photonics K.K.), a pathologist specializing in breast pathology annotated regions of invasive carcinoma, non-invasive carcinoma, normal ducts, vessels, tumor-infiltrating lymphocytes (TIL), and necrosis on the tissue sections. Areas where poor image quality made evaluation difficult were excluded from the annotations.

We used OpenAI’s GPT-4o for annotating the Visium tissue slices images as follows ^26^. For all samples with “hires” Visium images, we first removed the 150 pixels at the top, bottom, right and left sides, because the actual tissue slice is often located in the center of the image and not in its margins. The remaining cropped image was then cut into a 5-by-5 grid of rectangles of equal size. These 25 rectangles were passed on to GPT-4o using the Python openai package (version 1.36.0). We used the following prompt for samples with a known condition:

> *“This is a part of an H&E image of a <ORGANISM> tissue slice. The tissue is <TISSUE name>. The tissue is from a <CONDITION> individual. Describe any tissue features or signs of pathology in at most 100 characters. If there is not enough tissue to say anything, write ’empty’.”*

where <ORGANISM> is “human” or “mouse”, <TISSUE name> is one of the tissue names as included in the Uberon ontology, and <CONDITION> is one of the conditions included in the human DO. For samples lacking a condition annotation, the prompt was as follows:

> *“This is a part of an H&E image of a <ORGANISM> tissue slice. The tissue is <TISSUE name>. Describe any tissue features or signs of pathology in at most 100 characters. If there is not enough tissue to say anything, write ’empty’.”*

Responses were collected and are presented in the DeepSpaceDB database for each sample under section “Image annotation”.

### Database construction

The backend implementation of DeepSpaceDB employs Flask, a lightweight WSGI web application framework written in Python. Flask was selected due to its ease of use, adaptability, and wide range of extensions. While keeping a simple core that allowed us to use it for our specific architecture, the framework offers necessary routing, request handling, and API development tools. Modern JavaScript is used in the frontend’s construction, utilizing current web development techniques. The main objective of the implementation is to create a dynamic and responsive user experience for our analysis tools’ ease of use while adhering to clean design standards.

The main relational database is SQLite, which was selected due to its transaction capability, dependability, and serverless design. Sample-specific spatial transcriptomics data is kept in CSV format, as it can be easily transferred and backed up, and it is directly compatible with analysis tools. Plotly, a complete graphing package that provides interactive scientific charts and graphs, support for Python and JavaScript, and a variety of plot types appropriate for visualizing spatial transcriptomics data, is used to construct the visualization layer ^27^.

### Data availability

Data of all samples can be downloaded from the DeepSpaceDB website. Links to the original data source are also provided for each sample.

## Acknowledgement

The authors would like to thank Y. Harada for secretarial assistance. This work was supported by JST NBDC (Grant Number JPMJND2303, A.V.) and by an Office of Directors’ Research Grant provided by the Institute for Life and Medical Sciences of Kyoto University (A.Z. and A.V.). The funders had no role in study design, data collection and analysis, decision to publish, or preparation of the manuscript.

## Competing interests

The authors declare no competing interests exist in this study.

## Author contribution

A.V. conceived of the project, conducted data analysis, contributed to the database implementation, and wrote the manuscript. V.H. implemented the database and contributed to image data analysis. A.Z. contributed to data collection, data analysis, and database implementation. Y.H. conducted image annotation and interpretation. K.T. assisted in the database implementation. D.D. and S.K. contributed to the data analysis and database implementation. All authors contributed to critical revision of the manuscript.

## Notes

https://www.DeepSpaceDB.com

### Competing Interest Statement

The authors have declared no competing interest.

### Summary of Updates

Addition of new section "Analysis of uploaded samples" and minor revisions.

## References

1. Marx, V. Method of the Year: spatially resolved transcriptomics. Nat Methods 18, 9–14 (2021).

2. Ståhl, P. L. et al. Visualization and analysis of gene expression in tissue sections by spatial transcriptomics. Science (1979) 353, 78–82 (2016).

3. Vickovic, S. et al. High-definition spatial transcriptomics for in situ tissue profiling. Nat Methods 16, 987–990 (2019).

4. Fan, Z., Chen, R. & Chen, X. SpatialDB: A database for spatially resolved transcriptomes. Nucleic Acids Res 48, D233–D237 (2020).

5. Yuan, Z. et al. SODB facilitates comprehensive exploration of spatial omics data. Nat Methods 20, 387–399 (2023).

6. Xu, Z. et al. STOmicsDB : a comprehensive database for spatial transcriptomics data sharing , analysis and visualization. Nucleic Acids Res 1053–1061 (2024).

7. Li, Y., et al. SOAR elucidates disease mechanisms and empowers drug discovery through spatial transcriptomics. *bioRxiv* 2022.04.17.488596 (2023).

8. Bhuva, D. D. et al. Library size confounds biology in spatial transcriptomics data. Genome Biol 25, 1–10 (2024).

9. Janesick, A. et al. High resolution mapping of the tumor microenvironment using integrated single-cell, spatial and in situ analysis. Nat Commun 14, (2023).

10. Lu, M. Y. et al. A Multimodal Generative AI Copilot for Human Pathology. Nature (2024) doi:10.1038/s41586-024-07618-3.

11. Vandenbon, A. & Diez, D. A clustering-independent method for finding differentially expressed genes in single-cell transcriptome data. Nat Commun 11, 1–10 (2020).

12. Vandenbon, A. & Diez, D. A universal tool for predicting differentially active features in single-cell and spatial genomics data. Sci Rep 13, 1–14 (2023).

13. Cable, D. M. et al. Robust decomposition of cell type mixtures in spatial transcriptomics. Nat Biotechnol (2021) doi:10.1038/s41587-021-00830-w.

14. Lebrigand, K. et al. The spatial landscape of gene expression isoforms in tissue sections. Nucleic Acids Res 51, e47–e47 (2023).

15. Castranio, E. L. et al. Microglial INPP5D limits plaque formation and glial reactivity in the PSAPP mouse model of Alzheimer’s disease. Alzheimer’s and Dementia 19, 2239– 2252 (2023).

16. Haendel, M. A. et al. Unification of multi-species vertebrate anatomy ontologies for comparative biology in Uberon. J Biomed Semantics 5, 1–13 (2014).

17. Baron, J. A. et al. The DO-KB Knowledgebase: 20-year journey developing the disease open science ecosystem. Nucleic Acids Res 52, D1305–D1314 (2024).

18. Hao, Y. et al. Dictionary learning for integrative, multimodal and scalable single-cell analysis. Nat Biotechnol 42, 293–304 (2024).

19. Korsunsky, I. et al. Fast, sensitive and accurate integration of single-cell data with Harmony. Nat Methods 16, 1289–1296 (2019).

20. Diez, D. scmisc. Preprint at https://github.com/ddiez/scmisc.

21. Vandenbon, A. et al. Murine breast cancers disorganize the liver transcriptome in a zonated manner. Commun Biol 6, 1–12 (2023).

22. Dolgalev, I. msigdbr: MSigDB Gene Sets for Multiple Organisms in a Tidy Data Format. Preprint at (2022).

23. Megill, C. et al. cellxgene: a performant, scalable exploration platform for high dimensional sparse matrices. Preprint at 10.1101/2021.04.05.438318 (2021).

24. Han, Y. et al. TISCH2: expanded datasets and new tools for single-cell transcriptome analyses of the tumor microenvironment. Nucleic Acids Res 51, D1425–D1431 (2023).

25. Pagès, H., Carlson, M., Falcon, S. & Li, N. AnnotationDbi: Manipulation of SQLite-based annotations in Bioconductor. Preprint at https://bioconductor.org/packages/AnnotationDbi (2023).

26. OpenAI et al. GPT-4 Technical Report. (2023).

27. Plotly Technologies Inc. Plotly. Collaborative data science https://plot.ly (2015).

