## Supplementary Material for "DeepSpaceDB: a spatial transcriptomics atlas for interactive in-depth analysis of tissues and tissue microenvironments"

#### Supplementary Tables

| Feature | SODB | SOAR | STOmicsDB | DeepSpaceDB |
|---|---|---|---|---|
| Covers many platforms | yes | yes | yes | no |
| Includes quality indicators | no | no | yes | yes |
| Includes image data | no | yes | partly | yes |
| Includes image annotations | no | no | no | yes |
| Finding similar samples / overview of samples | no | no | no | yes |
| Gene expression visualization | yes | yes | yes | yes |
| Pathway activities | no | no | no | yes |
| Spot clustering | yes | yes | yes | yes |
| Cell type predictions | no | yes | yes | yes |
| Cell-cell interactions | no | no | yes | no |
| Interactive comparison of regions within a tissue slice | no | no | no | yes |
| Interactive comparison between slices | no | no | no | yes |
| Search for samples using query gene | no | no | yes | yes |
| Upload and analysis of own samples | no | no | no | yes |

Suppl. Table S1: Features included in DeepSpaceDB and three other spatial transcriptomics databases.

| Source name | URL | Number of samples |
|---|---|---|
| NCBI Gene Expression Omnibus | www.ncbi.nlm.nih.gov/geo/ | 1184 |
| EMBL-EBI | www.ebi.ac.uk/biostudies/ | 205 |
| Zenodo | zenodo.org | 141 |
| 10X Genomics | www.10xgenomics.com/datasets/ | 58 |
| Heart Cell Atlas | www.heartcellatlas.org/ | 42 |
| Lung Cell Atlas | www.lungcellatlas.org/ | 12 |
| Mendeley Data | data.mendeley.com | 8 |
| Reproductive Cell Atlas | www.reproductivecellatlas.org/ | 8 |
| Welcome Sanger Institute | treg-gut-niches.cellgeni.sanger.ac.uk | 8 |
| internal | - | 8 |

Suppl. Table S2: Sources of spatial transcriptomics samples using for DeepSpaceDB.

### Supplementary Figures

**Suppl. Figure S1: Tendencies of the samples included in DeepSpaceDB version 1.0**

**(A)** The cumulative number of Visium samples included in DeepSpaceDB version 1.0 in function of their publication data.

**(B-C)** Distribution of the number of reads per spot (X axis) versus the number of detected genes per spot (Y axis) for human (B) and mouse (C) samples. Colors indicate the number of spots per bin.

**(D-E)** Boxplots of the number of detected genes per spot (X axis) for each tissue of origin (Y axis) for human (D) and mouse (E) samples. There are considerable differences between the tissues, although the tendencies have to be interpreted carefully because differences can also be to some degree caused by differences in sequencing depts between studies.

**(F-G)** Boxplots showing the number of detected genes per spot (Y axis) in function of the number of immediate neighboring spots a spot has (X axis), for human (F) and mouse (G) samples. The number of immediate neighboring spots ranges from 0 (spots with no neighboring spots; i.e., isolated spots) up to 6 (spots with the maximum number of neighbors in the hexagonal grid of the Visium platform).

**(H-I)** The same as (F-G) but showing the distribution of the number of reads per spot in the Y axis, for human (H) and mouse (I) samples.

**Suppl. Figure S2: Embeddings of Visium pseudobulk samples and Visium spots in DeepSpaceDB**

**(A)** Embedding (tSNE plot) of mouse samples after processing to pseudobulk data. Colors indicate the tissue of origin. Prominent tissues are indicated.

**(B)** Embedding (tSNE plot) of human samples after processing to pseudobulk data. Colors indicate whether the sample was obtained from a patient suffering from cancer (blue) or not (red).

**(C)** Embedding (tSNE plot) of human spots. Colors indicate the 50 clusters obtained using k-means clustering.

**(D)** Embedding (tSNE plot) of mouse spots. Colors indicate the 50 clusters obtained using k-means clustering.

**Suppl. Figure S3: Properties of human spot clusters**

**(A,B)** Overlap between spot clusters and tissue annotations **(A)**, and between spot clusters and conditions annotations **(B)**. Overlap between sets of spots was estimated using the Jaccard index, which was converted to Z scores through randomizations (see Methods). Red colors indicate strong overlaps.

**(C)** Average scaled gene expression of selected differentially expressed genes of the 50 clusters. Annotations of clusters are indicated at the bottom of the figure.

**Suppl. Figure S4: Properties of mouse spot clusters**

**(A,B)** Overlap between spot clusters and tissue annotations **(A)**, and between spot clusters and conditions annotations **(B)**. Overlap between sets of spots was estimated using the Jaccard index, which was converted to Z scores through randomizations (see Methods). Red colors indicate strong overlaps.

**(C)** Average scaled gene expression of selected differentially expressed genes of the 50 clusters. Annotations of clusters are indicated at the bottom of the figure.

**Suppl. Figure S5: Examples of the distribution of annotated human spot clusters**

Each plot shows the distribution of spots within a 2D embedding (tSNE plot). This is the same embedding as shown in Suppl. Fig. S2C. To better reflect the density of spots, the space was divided into hexagonal bins, and the number of spots per bin is indicated by the intensity of the color (black: high, white: low). The red dotted lines show the rough contours of all spots in the 2D space as shown in panel **(A)**, to facilitate comparison of distributions between panels.

**(A)** Distribution of all human spots.

**(B-I)** A number of example annotations and the distribution of spots with these annotations.

The annotation is shown on top of each plot.

**Suppl. Figure S6: Examples of the distribution of annotated mouse spot clusters**

Each plot shows the distribution of spots within a 2D embedding (tSNE plot). This is the same embedding as shown in Suppl. Fig. S2D. To better reflect the density of spots, the space was divided into hexagonal bins, and the number of spots per bin is indicated by the intensity of the color (black: high, white: low). The red dotted lines show the rough contours of all spots in the 2D space as shown in panel **(A)**, to facilitate comparison of distributions between panels.

**(A)** Distribution of all mouse spots.

**(B-I)** A number of example annotations and the distribution of spots with these annotations. The annotation is shown on top of each plot.

**Suppl. Figure S7: Additional comparisons of gene expression between different regions within a sample**

**(A)** Scatterplot of the average gene expression in set 1 (X axis) and set 3 (Y axis). A number of genes with large differences is indicated.

**(B)** The spatial expression patterns of two selected genes are shown. *SLC12A2* has a higher expression in set 1 than in set 3. *FN1* has a higher expression in set 3 than in set 1.

**(C)** Scatterplot of the average gene expression in set 2 (X axis) and set 3 (Y axis). A number of genes with large differences is indicated.

**(D)** The spatial expression patterns of two selected genes are shown. *FASN* has a higher expression in set 2 than in set 3. *IGHA1* has a higher expression in set 3 than in set 2.

**Suppl. Figure S8: Summary of the Upload Center of DeepSpaceDB**

Users can upload Visium samples and process them in a similar way to the samples included in the DeepSpaceDB database. Users can give a job label, select the species and the image resolution of the uploaded image data. A human or mouse example sample can also be used. The data to upload consists of output files of the 10X Genomics Space Ranger software: 1) a barcode file, 2) a feature file, 3) a matrix file, 4) an image file, 5) a scalefactors file, and 6) a tissue position file. After uploading these data files, their content is checked on our server, and – if no problems are found – the data is submitted by our job scheduling software for processing. The status of the sample will be listed under “Recent jobs”. After the processing has been completed, the uploaded sample can be inpected on the server. Alternatively, the sample can be compared with other samples, the processed data can be downloaded, or the sample can be deleted. Uploaded samples are automatically deleted after some time. Each uploaded sample is accessible through a unique URL, which can be shared with collaborators. Data is not visible or accessible to others without the unique URL. Uploaded data is not included in the DeepSpaceDB database, and will not be collected or retained on our side.
