## Supplementary figures and images for "DeepSpaceDB: a spatial transcriptomics atlas for interactive in-depth analysis of tissues and tissue microenvironments"

### Supplementary Figures

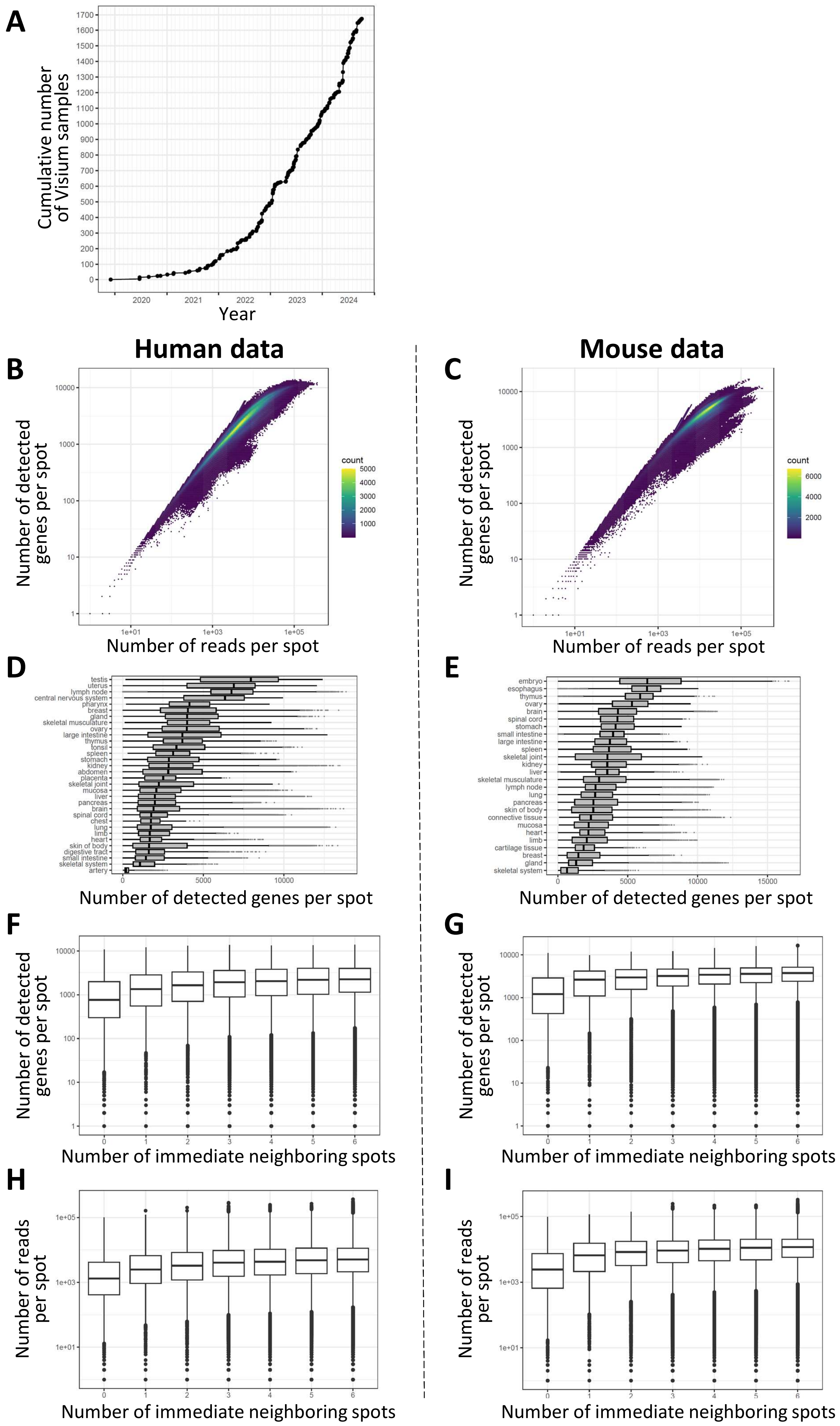

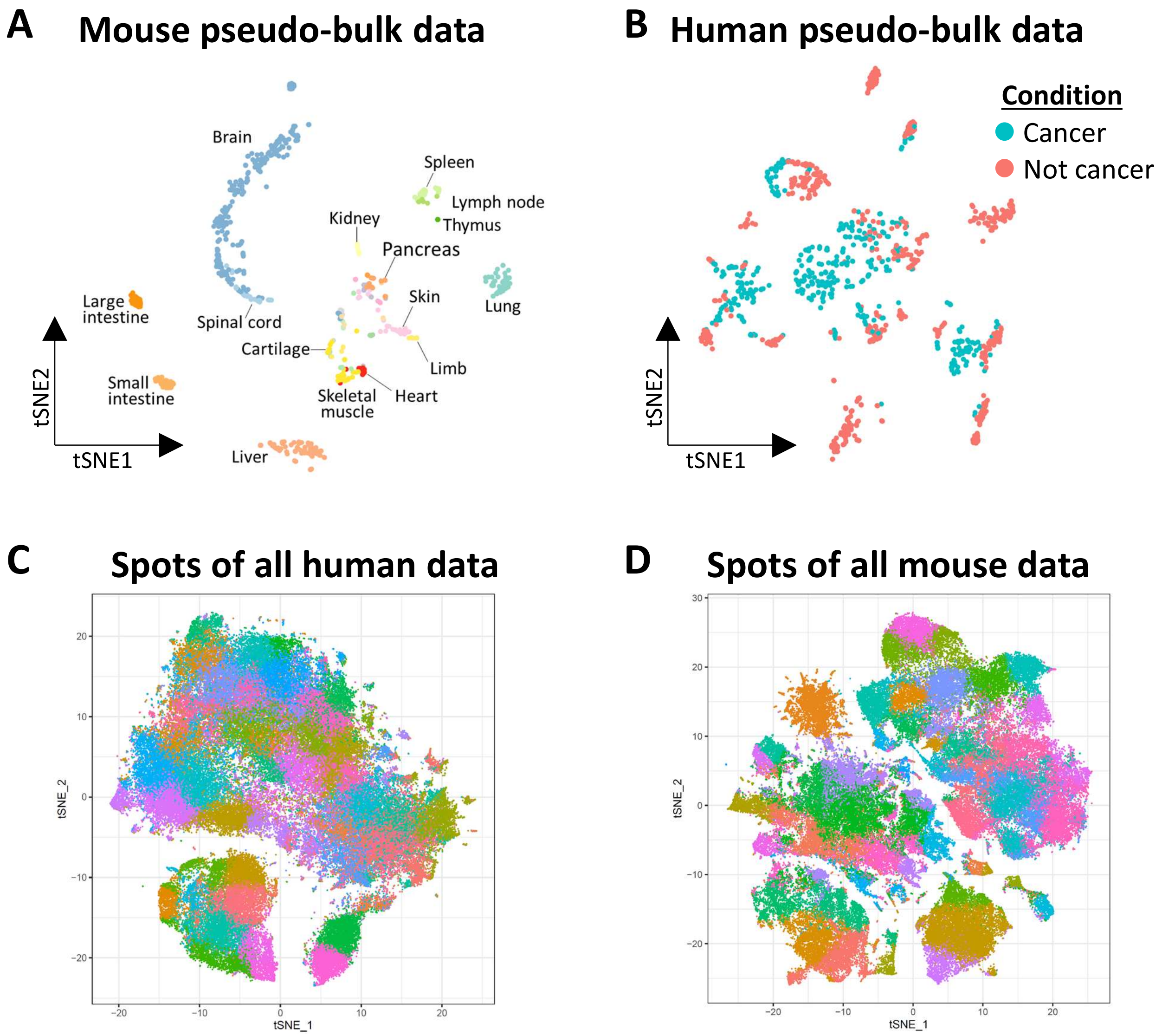

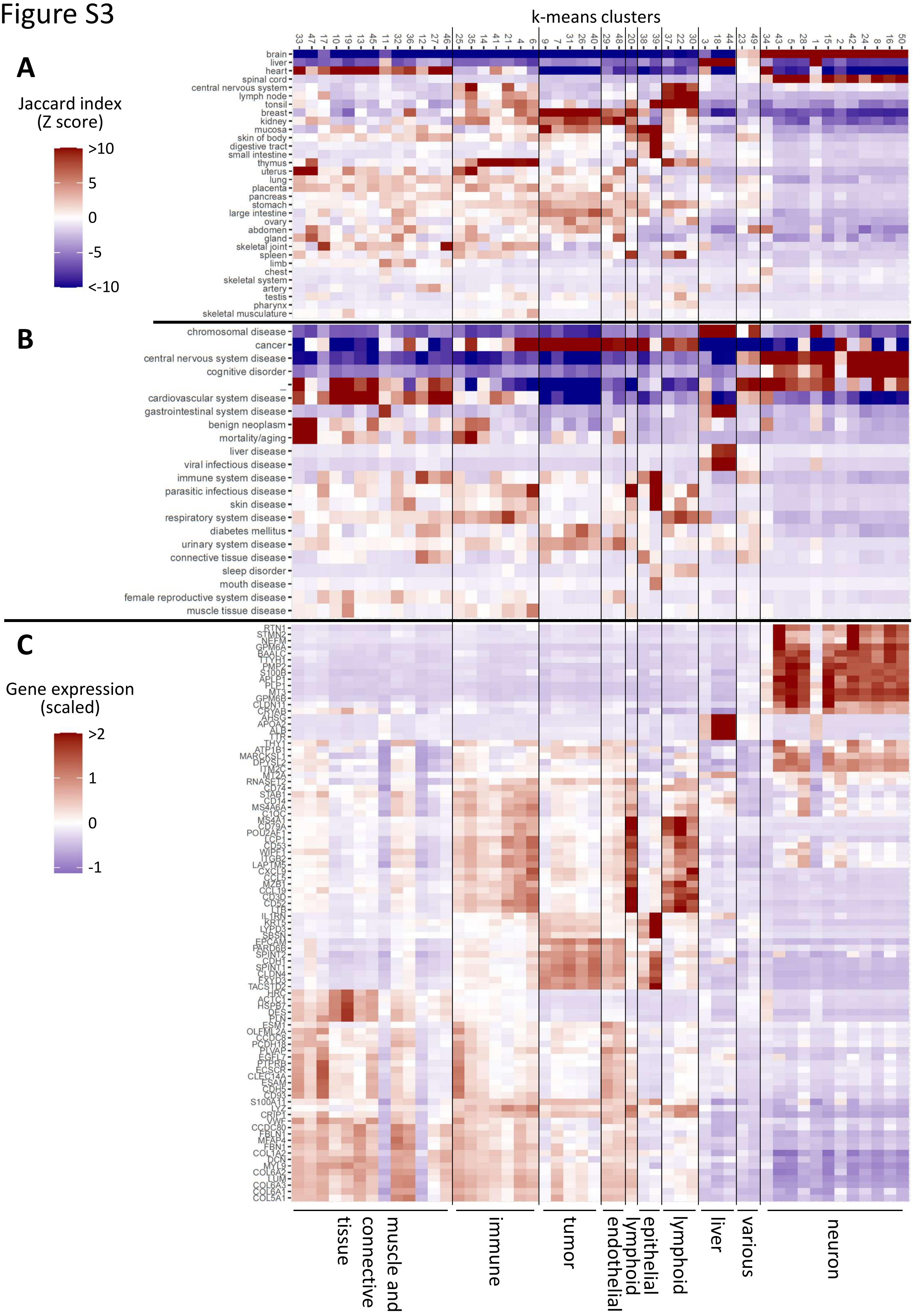

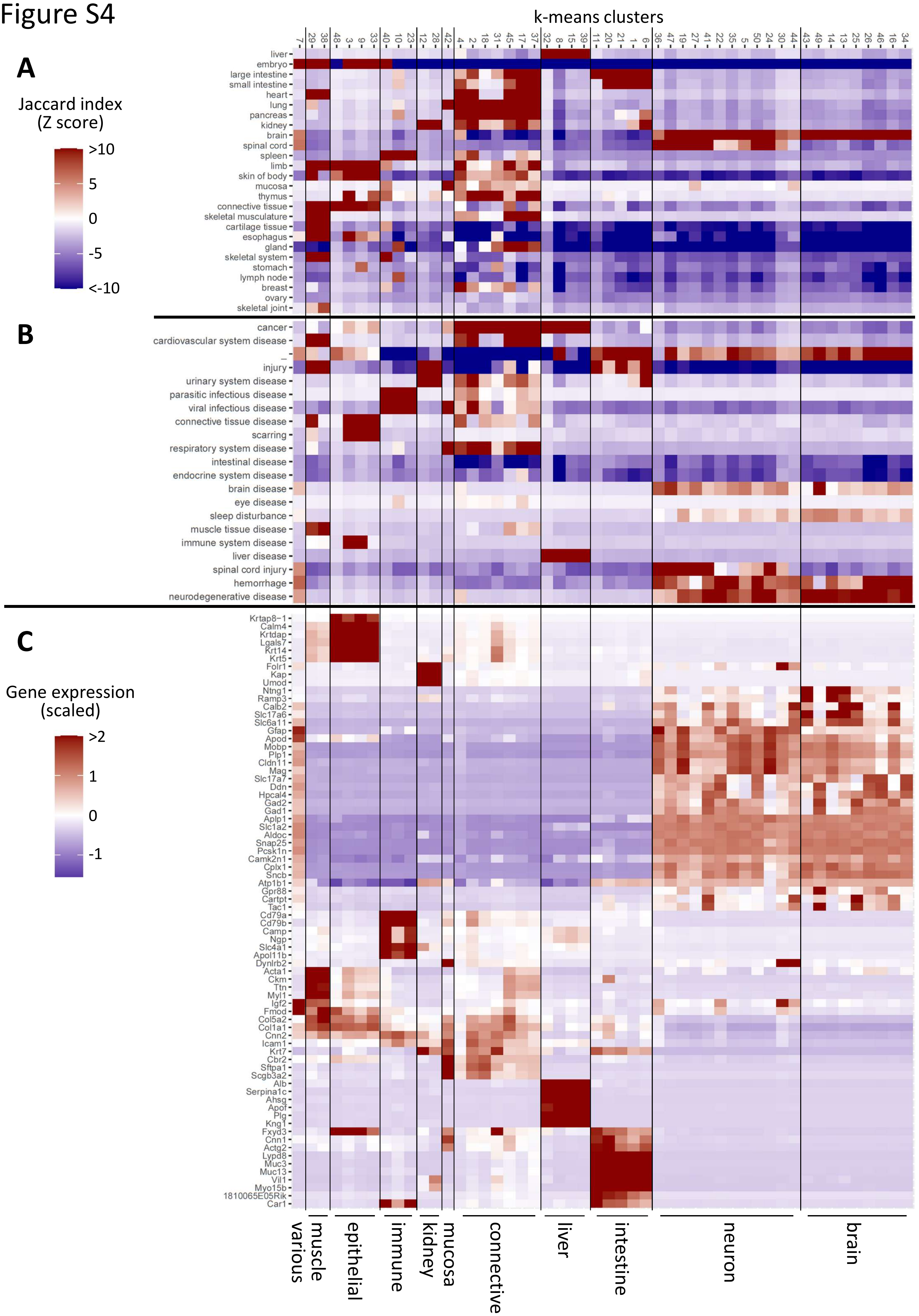

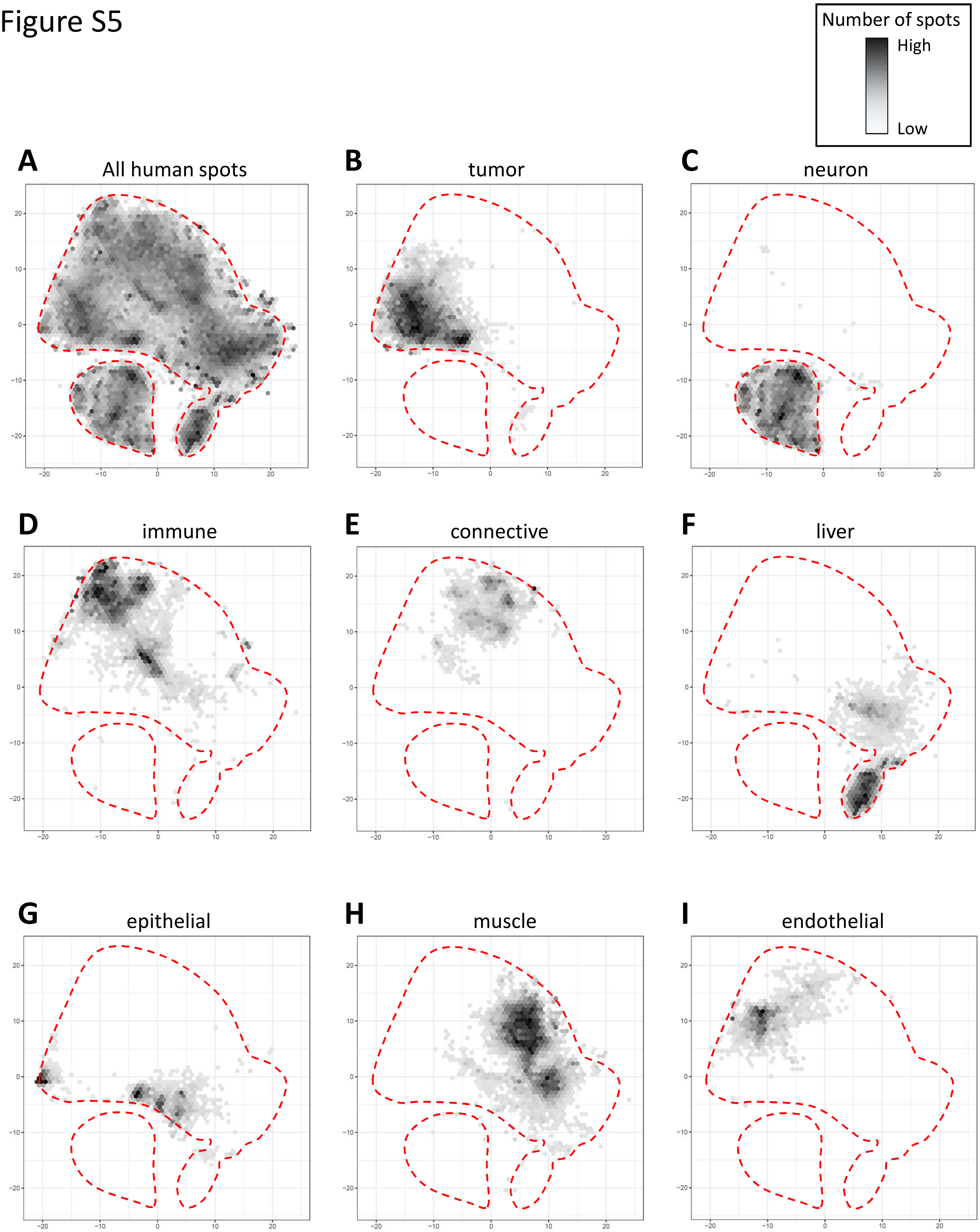

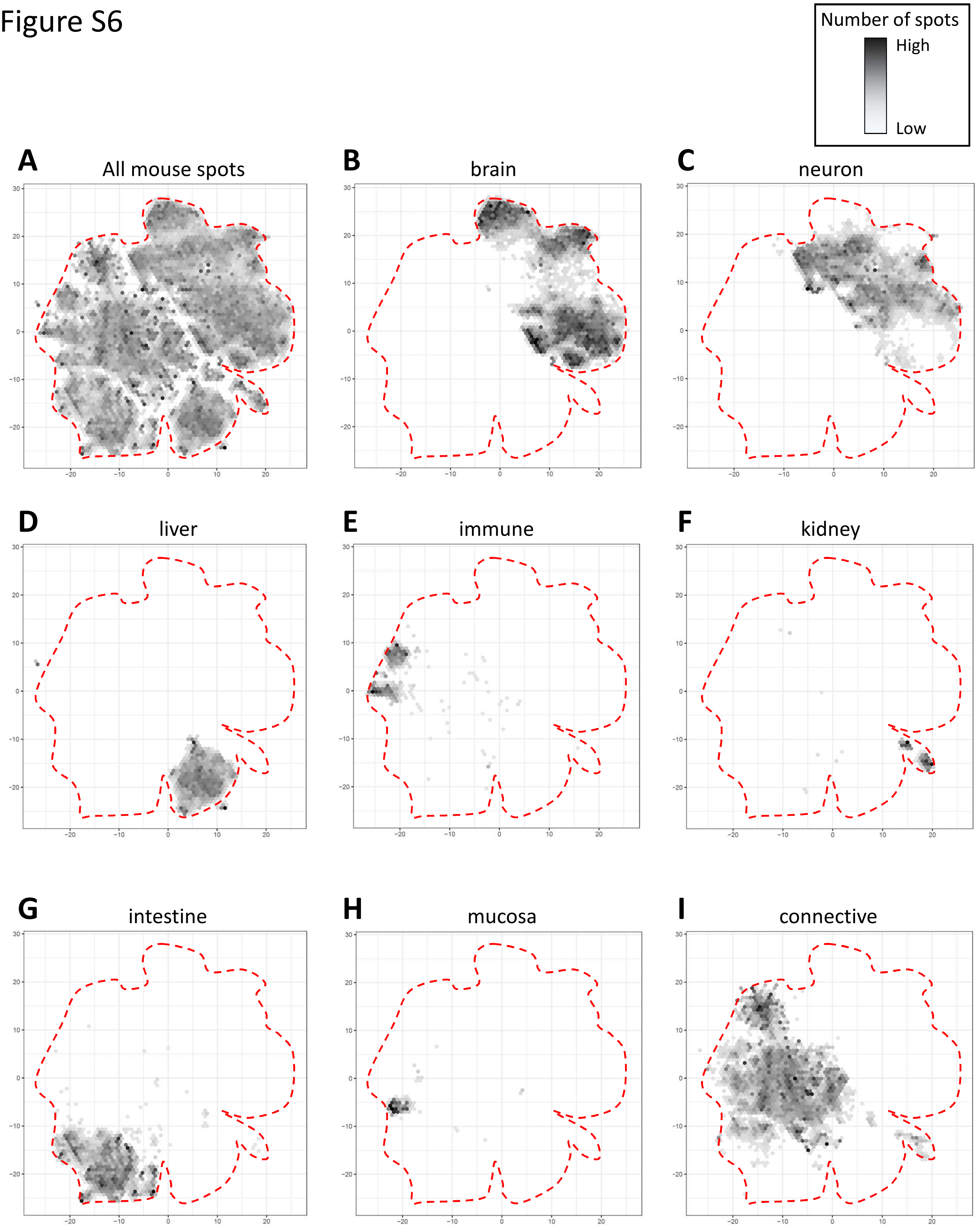

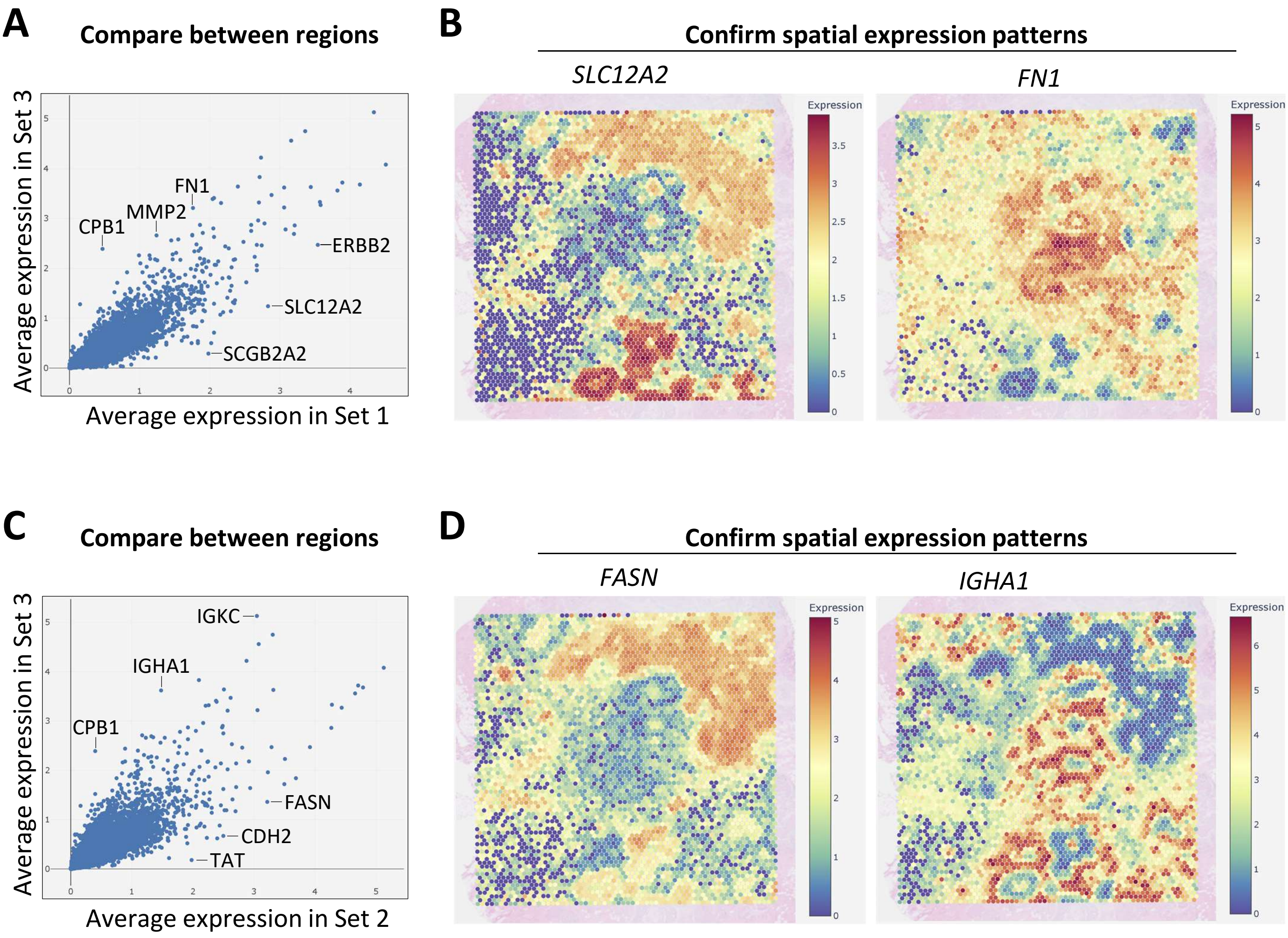

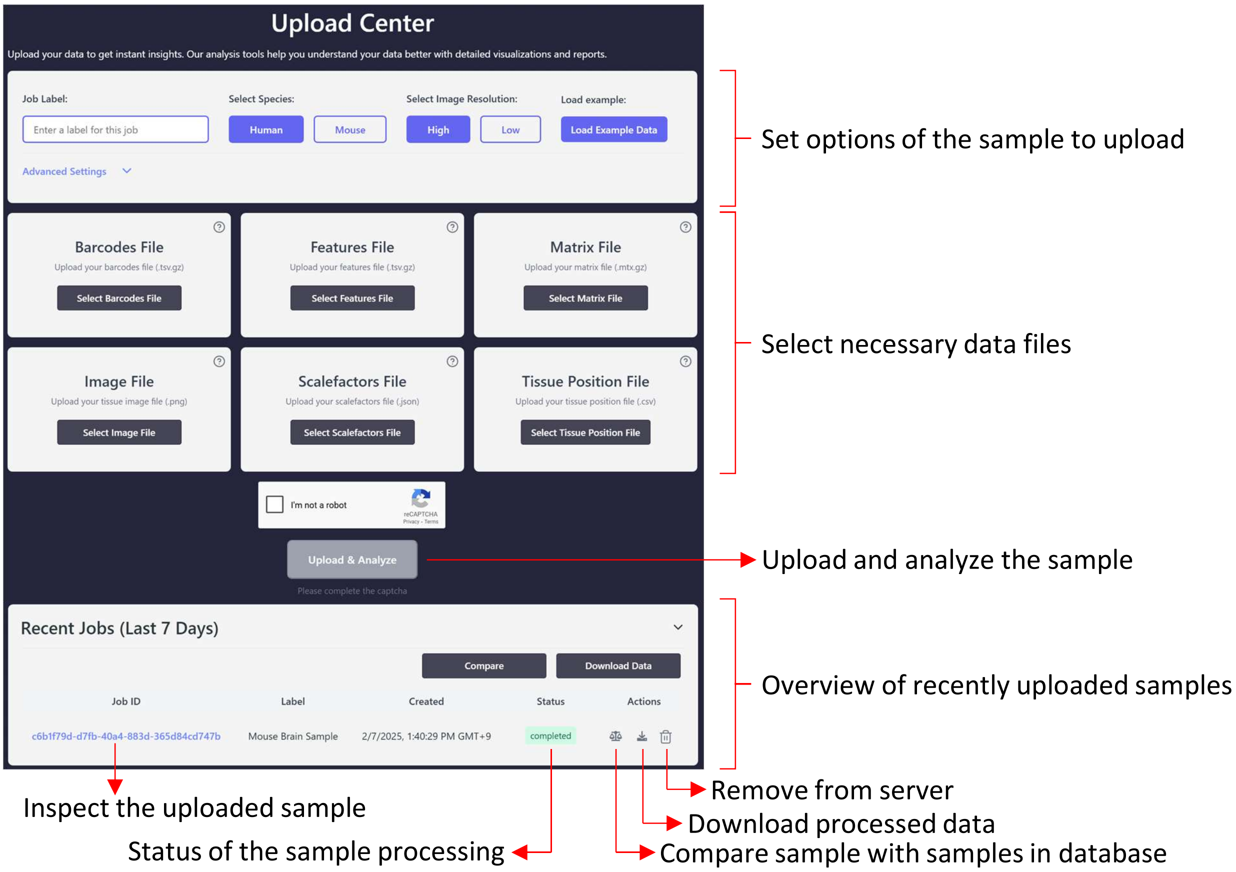
